# Mining single-cell transcriptomic data reveals distinct T-cell population in pediatric B-ALL and AML at diagnosis

**DOI:** 10.64898/2026.01.09.698676

**Authors:** Liqing Tian, Stephen Gottschalk

## Abstract

Immunotherapy represents a promising strategy to improve outcomes in pediatric leukemia; however, its efficacy remains considerably lower in acute myeloid leukemia (AML) compared with B-cell acute lymphoblastic leukemia (B-ALL). To characterize T-cell subsets in AML and B-ALL at diagnosis, we took advantage of existing data sets and performed an integrated single-cell RNA sequencing (scRNA-seq) analysis of T cells present in the bone marrow of pediatric patients with B-ALL (n=89) and AML (n=26), and healthy donors (n=9). In total, 47,610 T cells were analyzed, revealing 17 transcriptionally distinct subsets. Comparative analysis identified T-cell subsets distinguishing B-ALL from AML, such as proliferative, naïve CD4, and progenitor exhausted, and naïve or resting CD4 regulatory T cells. We identified a rare T-cell subset expressing hemoglobin genes, which was enriched in B-ALL and characterized by upregulation of heme metabolism and chronic hypoxia-associated pathways, and its abundance was associated with better outcomes. Collectively, our findings delineate the transcriptional and functional heterogeneity of T cells in pediatric B-ALL and AML and provide insights that may inform future T-cell-based immunotherapeutic strategies.

## INTRODUCTION

Immunotherapy has emerged as an attractive approach to improve outcomes for pediatric patients with B-ALL and AML.^1^ T-cell-based immunotherapy approaches such as bispecific T-cell engagers and chimeric antigen receptor (CAR) T cells harness the patient’s own immune system to target tumor cells. While these strategies have achieved remarkable success for B-ALL, their efficacy in AML remains limited, reflecting fundamental differences in tumor biology and the immune microenvironment between these leukemia subtypes.^2^

Increasing evidence highlights the critical role of the immune microenvironment, particularly T cells, in shaping disease progression, treatment response, and relapse dynamics in leukemia.^3,4^ However, the composition and functional diversity of T cells within the pediatric leukemic bone marrow remain poorly understood. Characterizing these T-cell populations is critical for elucidating mechanisms of immune evasion and for optimizing immunotherapeutic interventions. Mining available scRNA-seq data sets that were primarily generated to study leukemia blasts presents a unique opportunity to dissect T-cell subsets in the bone marrow of pediatric patients with B-ALL and AML.

In this study, we performed an integrated single-cell transcriptomic analysis of T cells derived from the bone marrow of a large cohort of pediatric patients with B-ALL and AML, as well as healthy donors. Our findings delineate the transcriptional landscape of T cells within the pediatric leukemic bone marrow and provide insights that may guide the development of effective T cell-based immunotherapies for childhood leukemia.

## METHODS

scRNA-seq data from patients with B-ALL (89 samples^5^, SCPCP000008) and AML (26 samples, SCPCP000007) were obtained from the Single-cell Pediatric Cancer Atlas (ScPCA) portal.^6^ Most samples were derived from bone marrow (3 from peripheral blood) at initial diagnosis (7 from recurrence cases). scRNA-seq data from healthy pediatric or young adult bone marrow donors (9 samples) were collected from GSE132509^7^, GSE154109^8^, and SCPCP000007 (ScPCA). Details of the discovery dataset are provided in **supplemental Table 1**. An independent validation cohort was analyzed, consisting of scRNA-seq data from 7 pediatric B-ALL and 7 pediatric AML samples, all derived from bone marrow samples at diagnosis (GSE154109^8^, details in **supplemental Table 2**). Additionally, a scRNA-seq dataset of peripheral blood mononuclear cells (PBMCs) from healthy donors (10x Genomics) was included.

All samples in this study contained at least 800 cells after low-quality cells were filtered out. T cells were extracted from each sample and pooled for analysis across B-ALL, AML, and healthy donor samples. T-cell subsets were identified and annotated using established marker genes^9^ and details are provided in the Supplemental Material section. To ensure robust statistical comparison of T cell subset fractions, only samples containing at least 50 T cells were included, resulting in 85 B-ALL, 20 AML, and 9 healthy donor bone marrow samples in the discovery dataset.

For pathway enrichment analysis, average gene expression per cluster was computed using the Seurat AverageExpression function^10^, and pathway enrichment scores were computed using the GSVA package^11^ with hallmark and curated gene sets from MSigDB^12^. Marker genes for each T cell cluster were identified using Seurat FindAllMarkers^10^ (adjusted p < 0.05, ranked by average log2 fold change). The Seurat AddModuleScore function^10^ was used to compute gene signature scores for each cell. Additional methods are provided in the Supplemental Material. The processed datasets used in this analysis are available on GitHub (https://github.com/liqingti/T_cells_in_pediatric_B-ALL_AML_healthydonors).

## RESULTS AND DISCUSSION

We analyzed T cells from the bone marrow of healthy donors and pediatric patients with B-ALL or AML, yielding 47,610 T cells from 124 samples. Unsupervised clustering identified 17 T-cell subsets, including CD4 naïve, CD4 regulatory (Treg), CD8 naïve, CD8 effector, CD8 exhausted, mixed CD4/CD8 subsets (T_mixed) such as proliferative T, γδ T (Tgd) or mucosal-associated invariant T (MAIT) cells (**Figure 1A–C**). Naïve CD4 T cells were the most abundant T-cell subset, comprising the majority of CD4 T cells (**Figure 1C**).

**Figure 1.**
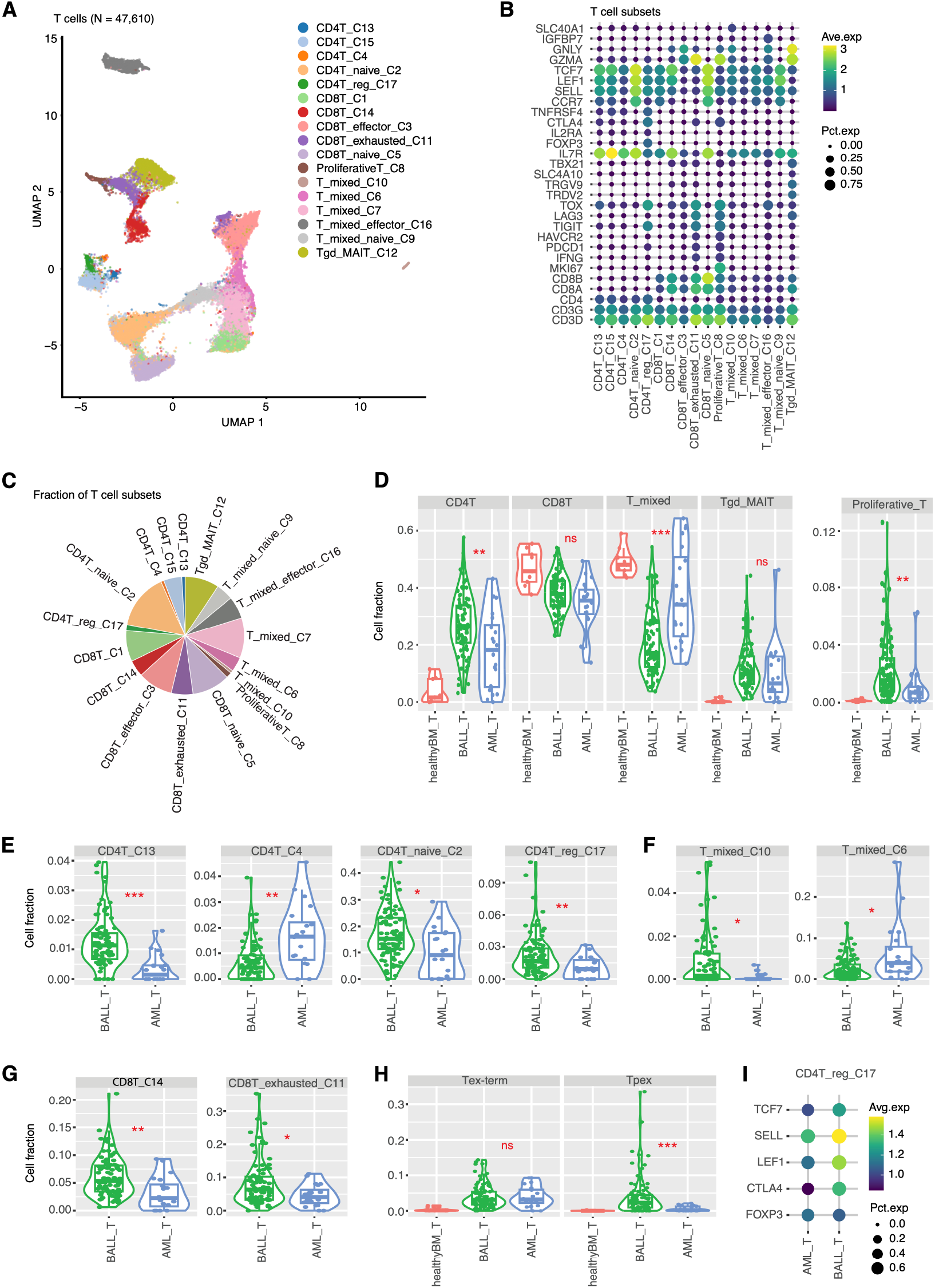
Characteristics of T-cell subsets in patients with B-ALL or AML, and healthy bone marrow donors. (**A**) UMAP view of 17 distinct T-cell clusters. (**B**) Marker gene expression across defined T-cell clusters. Bubble size indicates the proportion of cells expressing each gene, and color represents the average expression level. (**C**) Fraction distribution of T-cell subsets. (**D**) Comparison of major T-cell subset fractions. (**E-G**) T-cell fraction comparisons showing significant differences among (**E**) CD4 T, (**F**) T mixed, and (**G**) CD8 T subsets. (**H**) Comparison of cell fractions between Tex-term and Tpex cells within the CD8T_exhausted_C11 subset. (**I**) Gene expression in the CD4T_reg_C17 subset across AML and B-ALL. Statistical significance was determined using two-sided Wilcoxon rank-sum tests, with P values adjusted for multiple testing by the false discovery rate (FDR) method. FDR-adjusted P values: *<0.05; **<0.01; ***<0.001. Boxes indicate interquartile ranges with medians. Samples with fewer than 50 T cells were excluded from analyses in panels D-H.

A comparative analysis of the fractions of major T-cell subsets (CD4, CD8, T_mixed, Tgd_MAIT, and proliferative T) showed significant differences for CD4, T_mixed, and proliferative T-cell subsets between B-ALL and AML (**Figure 1D**). Of the 17 identified T-cell subsets, eight had significantly different fractions between B-ALL and AML (**Figure 1E–G, supplemental Figure 1**). Proliferative T cells accounted for 1.56% of all T cells (**Figure 1C**) and were almost absent in healthy donor bone marrow (**Figure 1D**). Notably, B-ALL had a higher fraction of proliferative T cells compared with AML. Given the previously established association between proliferative T cells and favorable outcomes in solid tumors,^13,14^ this analysis suggests that the presence of proliferative T cells could be a prognostic indicator in pediatric leukemia.

CD4 T cells were significantly more abundant in B-ALL than AML samples, predominantly comprising naïve CD4 T cells (**Figure 1D–E**). Exhausted CD8 T cells were almost absent in healthy donor bone marrow (**supplemental Figure 1**), and more abundant in B-ALL than AML samples (**Figure 1G**). However, this was due to a higher fractions of progenitor exhausted T (Tpex) and not terminally exhausted T (Tex-term) cells (**Figure 1H**), with the former being associated with better therapeutic responses in preclinical models as well as clinical studies.^15^ Tregs were almost absent in healthy donor bone marrow (**supplemental Figure 1**), consistent with one previous study.^16^ Tregs were more abundant in B-ALL than AML (**Figure 1E**). Although the expression of the key Treg marker FOXP3 was comparable between the two groups, naïve T-cell markers (LEF1, SELL, TCF7) and the inhibitory marker CTLA4 were expressed at higher levels in B-ALL samples (**Figure 1I**). These finding suggests that B-ALL samples harbor more naïve or resting Treg cells than AML.

A rare subset, T_mixed_C10, was clearly separated from other T-cell subsets on UMAP (**Figure 1A**) and was more frequent in B-ALL than AML (**Figure 1F**). This subset was also enriched in standard-risk compared to high-risk B-ALL (**Figure 2A**), suggesting a potential protective role in pediatric leukemia. Pathway analysis revealed upregulation of heme metabolism and reactive oxygen species (ROS) pathways (**Figure 2B**). Hemoglobin genes (α-, β-, γ-, δ-globin) were among the top markers distinguishing T_mixed_C10 cells from other T cell subsets (**Figure 2C**). Validation confirmed that T_mixed_C10 subset was not an artifact since (1) the detected gene numbers were comparable to other subsets (**supplemental Figure 2A**), (2) doublet scores were comparable to other subsets (**supplemental Figure 2B**), and (3) the expression of hemoglobin genes was similar between CD3 high and CD3 low cells in the subset (**supplemental Figure 2C**) supporting no erythroid contamination.

**Figure 2.**
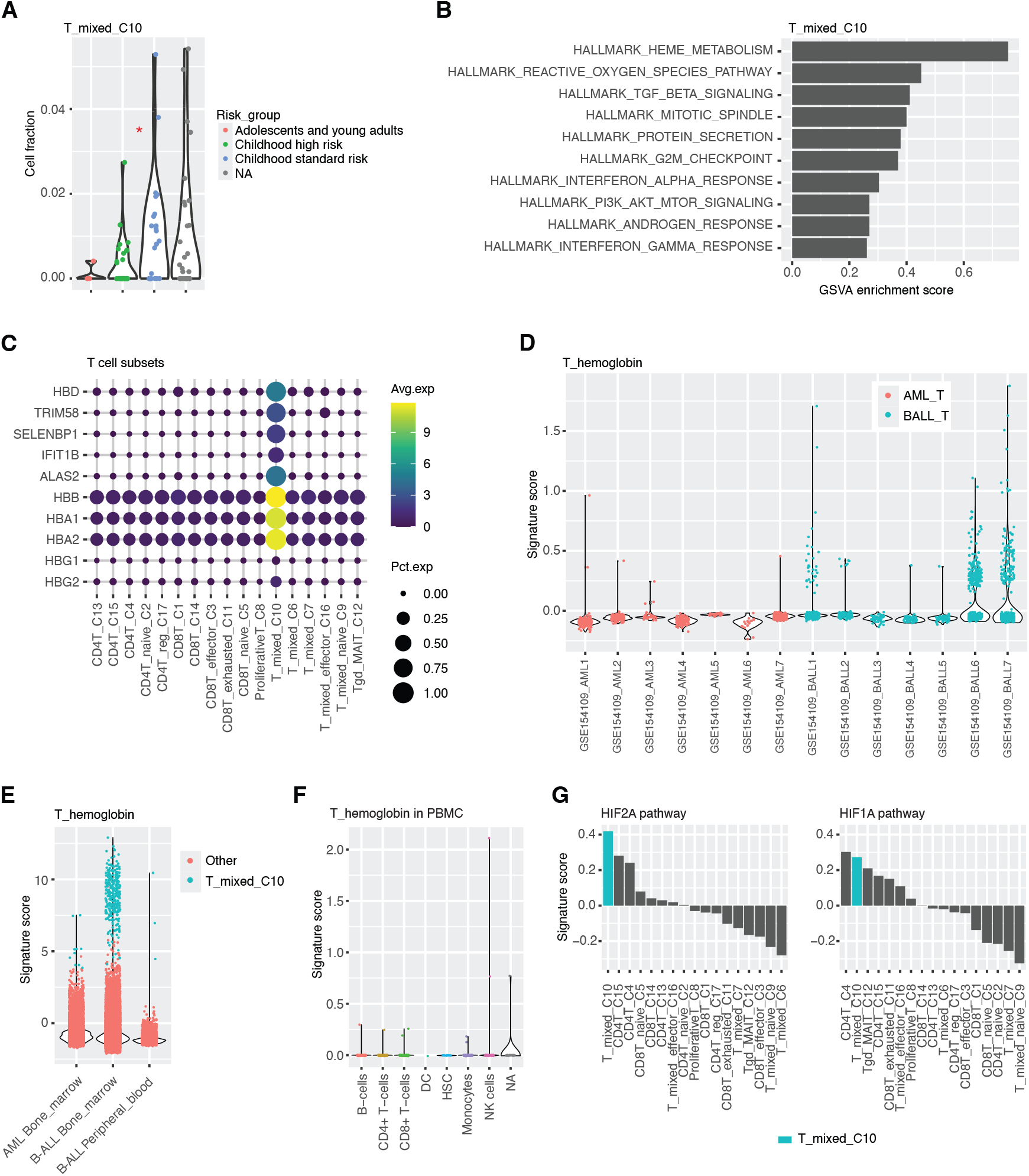
Characteristics of hemoglobin-expressing T cells. (**A**) Comparison of cell fractions among B-ALL risk groups within the T_mixed_C10 subset. (**B**) Top enriched hallmark pathways in T_mixed_C10 cells. (**C**) Top marker genes distinguishing T_mixed_C10 cells from other T-cell subsets. (**D**) T_hemoglobin (hemoglobin-expressing T-cell) signature scores in an independent cohort of patients with B-ALL and AML. (**E**) T_hemoglobin signature scores in bone marrow and peripheral blood samples from patients with B-ALL and AML. (**F**) T_hemoglobin signature scores in PBMCs from healthy donor. (**G**) HIF2A and HIF1A pathway signature scores across T-cell subsets. Statistical significance was determined using two-sided Wilcoxon rank-sum tests. P values: *<0.05. Samples with fewer than 50 T cells were excluded from analysis in panel A.

In the validation cohort, T_hemoglobin signature high T cells were observed almost exclusively in B-ALL (**Figure 2D**), consistent with the discovery cohort (**Figure 2E**). Within B-ALL of the discovery cohort, these cells were predominantly found in bone marrow, but rarely in peripheral blood (**Figure 2E**). Likewise, healthy donor PBMC data revealed very few peripheral blood T cells with high T_hemoglobin signature score (**Figure 2F**). Pathway analysis revealed that the chronic hypoxia-associated HIF2 signaling pathway^17^ was markedly elevated compared to acute HIF1 signaling pathway (**Figure 2G**) in this subset. These findings suggest that this rare, bone marrow– restricted T-cell subset arises in response to chronic hypoxic stress within the B-ALL bone marrow microenvironment. A recent study supports the notion that T cells can express hemoglobin genes, potentially contributing to redox regulation and mitochondrial function.^18^ Moreover, heme supplementation has been shown to enhance effector T- and CAR T-cell functions while preventing T-cell exhaustion.^19^ Collectively, these observations point to a potential role for hemoglobin-expressing T cells in antitumor immunity in pediatric B-ALL.

In conclusion, our study demonstrates significant differences of T-cell subsets at diagnosis in the bone marrow between pediatric AML and B-ALL. The results suggest that the endogenous immune system responds differently to AML and B-ALL blasts and future studies are needed to further delineate underlying mechanisms. In addition, our study highlights the power of reanalyzing existing datasets to investigate immune cells that were initially generated to study malignant cells.

## ACKNOWLEDGEMENTS

This work was supported by the American Lebanese Syrian Associated Charities (ALSAC) to SG.

## AUTHORSHIP CONTRIBUTIONS

LT and SG conceptualized the study. LT performed the analysis and created the figures, and SG provided funding. LT and SG wrote and edited the manuscript.

