## Supplemental Figures for "Mining single-cell transcriptomic data reveals distinct T-cell population in pediatric B-ALL and AML at diagnosis"

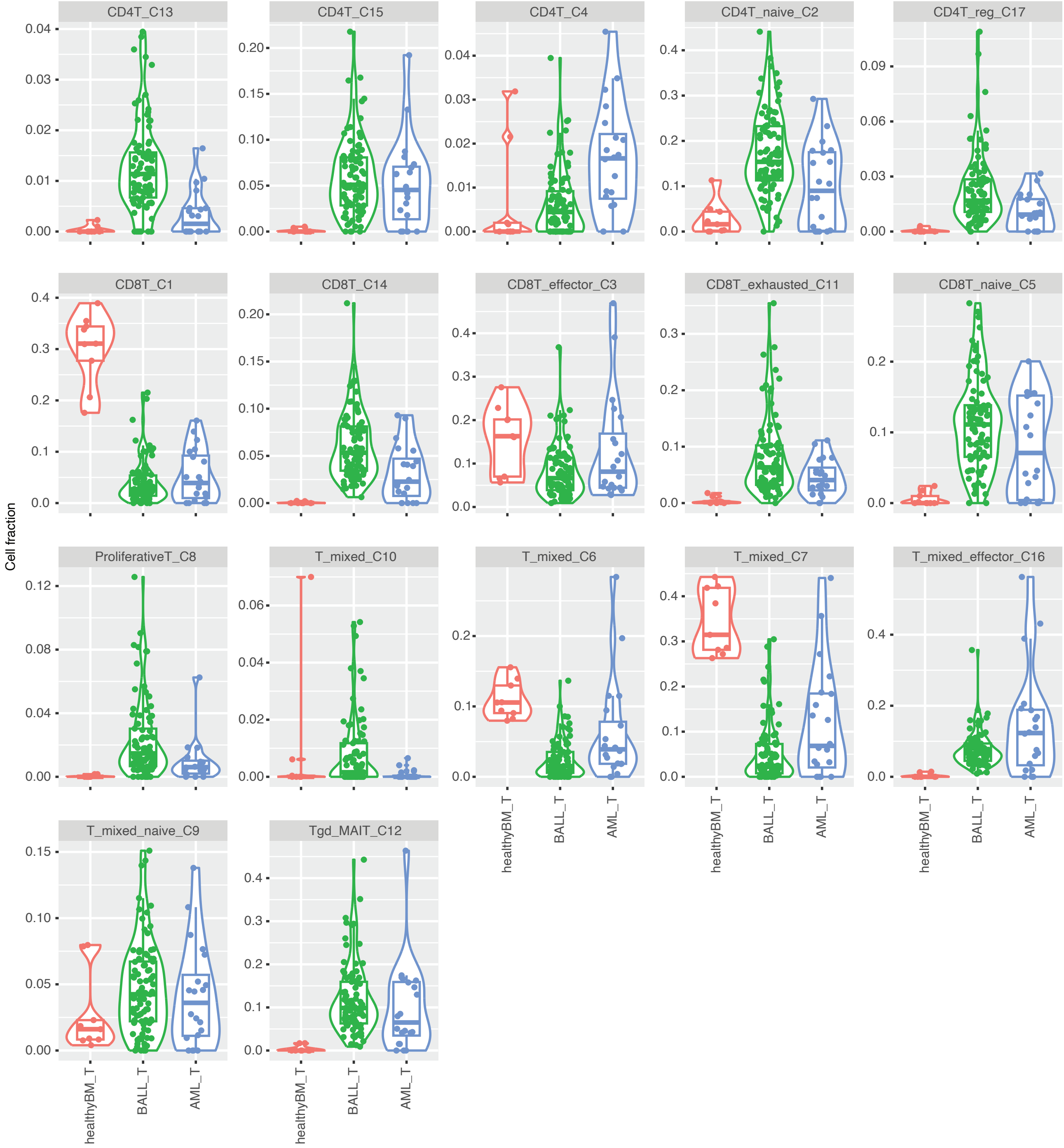


**Figure S1. Comparison of all T-cell subset fractions among B-ALL, AML and healthy bone marrow samples.** For detail see text.

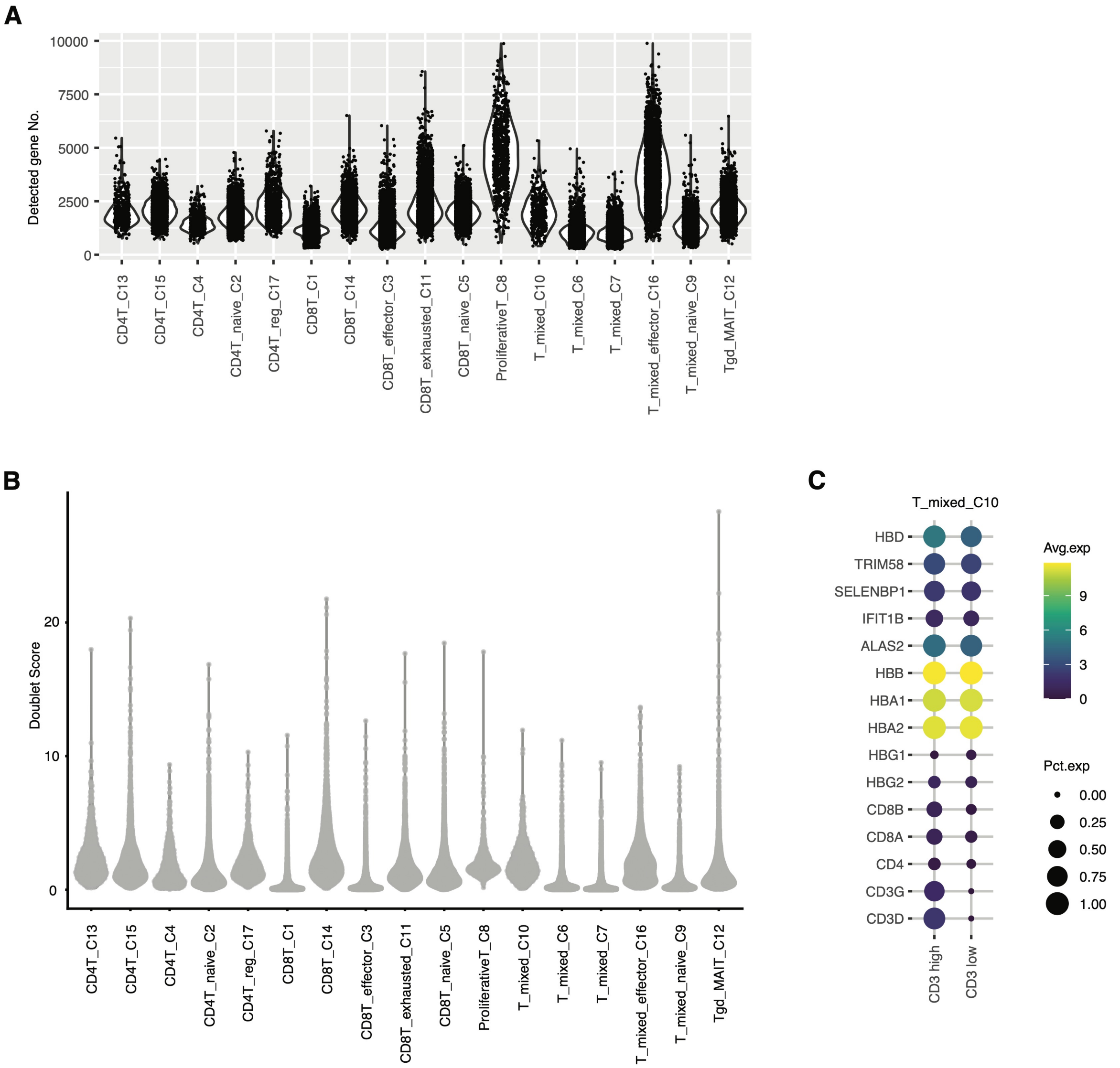


**Figure S2. Validation that T-cell subtype T_mixed_C10 is not an artifact.** (**A**) Distribution of detected gene number per cell across T-cell subtypes. (**B**) Distribution of doublet score per cell across T-cell subtype. (**C**) Marker gene expression in CD3 high and CD3 low cells (defined in supplemental methods) within the T_mixed_C10 subtype.
