## Supplemental Methods for "Mining single-cell transcriptomic data reveals distinct T-cell population in pediatric B-ALL and AML at diagnosis"

#### Supplemental Methods
#### for

### Mining single-cell transcriptomic data reveals distinct T-cell population in pediatric B-ALL and AML at diagnosis

Liqing Tian[1], Stephen Gottschalk[1]

[1]Department of Bone Marrow Transplantation and Cellular Therapy, St. Jude Children's Research Hospital, Memphis, TN 38105, USA

**scRNA-seq data processing for B-ALL, AML and healthy bone marrow samples**
Processed rds files and sample information for 89 B-ALL samples[1] were obtained from ScPCA portal (dataset: SCPCP000008)[2]. Data for 26 AML samples and two young adult healthy bone marrow samples were obtained from the same portal (dataset: SCPCP000007).

The ScPCA portal used a standardized pipeline to preprocess scRNA-seq data across all samples, and 3 cell-type annotation labels ("singler", "cellassign", "consensus") were assigned using automated methods. We selected cells annotated with T cell or natural killer (NK) cell by any of the 3 labels as processed cells for the following analysis.

Raw scRNA-seq data from GSE132509[3] and GSE154109[4] were also downloaded. For each sample, low-quality cells with high mitochondrial gene expression or fewer than 300 detected genes were filtered out. All remaining cells defined as processed cells were used for the following analysis. Gene expression was normalized using quickCluster and computeSumFactors (from scran package[5]) and logNormCounts (from scuttle package[6]).

The intersection of gene lists among the ScPCA, GSE132509 and GSE154109 datasets was used to construct the unified scRNA-seq data matrix for the samples from the 3 datasets. For each discovery cohort (B-ALL, AML, and healthy bone marrow samples), the processed cells were collected and run the following process separately. The top 2000 highly variable genes were selected to PCA computing using modelGeneVar, getTopHVGs, multiBatchPCA (from scran). Harmony algorithm (harmony package[7]) was used to integrating multiple samples. Batch correction and sample integration were performed using the harmony algorithm, and harmony PCA was used to generate UMAP view (plotReducedDim from scater[6]) and perform unsupervised clustering (clusterRows from bluster[8]). T cell clusters showing low T cell marker (CD3D, CD3G) expression or high NK markers (NCAM1, KLRB1, KLRD1, KLRF1, FCGR3A) - visualized using plotDots (scater) - were removed. Two rounds of filtering were performed to obtain pure T cells.

All purified T cells from the 3 cohorts were collected and reprocesses (highly variable genes selection, PCA, harmony PCA, UMAP, clustering, and marker gene expression visualization). The same workflow was applied to the validation B-ALL and AML cohort, with one round of T cell purification.

**scRNA-seq data processing for healthy donor peripheral blood mononuclear cells (PBMCs)**
The filtered matrix data was downloaded from 10X Genomics PBMC4k dataset (https://support.10xgenomics.com/single-cell-gene-expression/datasets/2.1.0/pbmc4k). As described above, low quality cells with high mitochondrial gene expression or fewer than 300 detected genes were filtered out. Cell types were annotated using SingleR (pruned.labels), with reference data from the BlueprintEncodeData function in the celldex package[9].

**T-cell subset annotation**
T-cell subsets were identified and annotated using established marker genes from pan-cancer T cell atlas[10]. Proliferative T cells were characterized by MKI67 expression. Tgd or MAIT cells were defined by high expression of TRDV2, TRGV9, SLC4A10 or TBX21. Exhausted T cells were defined by high expression of PDCD1 (PD-1), HAVCR2 (TIM-3), TIGIT, LAG3 and TOX. Treg cells were defined by low IL7R (CD127) expression, and high expression of FOXP3, IL2RA (CD25), CTLA4 and TNFRSF4. Naïve T cells were defined by high expression of CCR7, SELL, LEF1 and TCF7 (TCF-1). Effector T cells were defined by high expression of GZMA and GNLY. Figure 1B shows the marker gene expression across T cell subsets.

**Classification of Tpex and Tex-term cells**
Cells from the CD8T_exhausted_C11 subset were reprocesses (highly variable genes selection, PCA, harmony PCA, and clustering), yielding 17 subclusters. Based on expression of established Tpex marker genes (TCF7, CD27, CD28, CD38, SELL, CXCR3, GZMK)[10] subclusters 2, 4, 6, 12, 13, 14, 15, 16, and 17 were classified as Tpex cells, while the remaining subcluster cells were defined as Tex-term cells (**supplemental methods Figure 1**).

**CD3 high and CD3 low cells within the T_mixed_C10 subset**
To determine whether hemoglobin-expressing T cells in the subset were due to contamination of erythroid lineage cells, cell in the subset were divided into two groups. CD3 low group: cells lacking expression of both CD3D or CD3G (logcount = 0), and CD3 high group: all other cells.

**Doublet score**
Doublet scores were calculated using the computeDoubletDensity function (scDblFinder package[11]) per cell with harmony PCA as input.

**Gene signature score**
The T_hemoglobin signature was defined as gene set (HBA1, HBA2, HBB, HBD) based on marker gene expression.

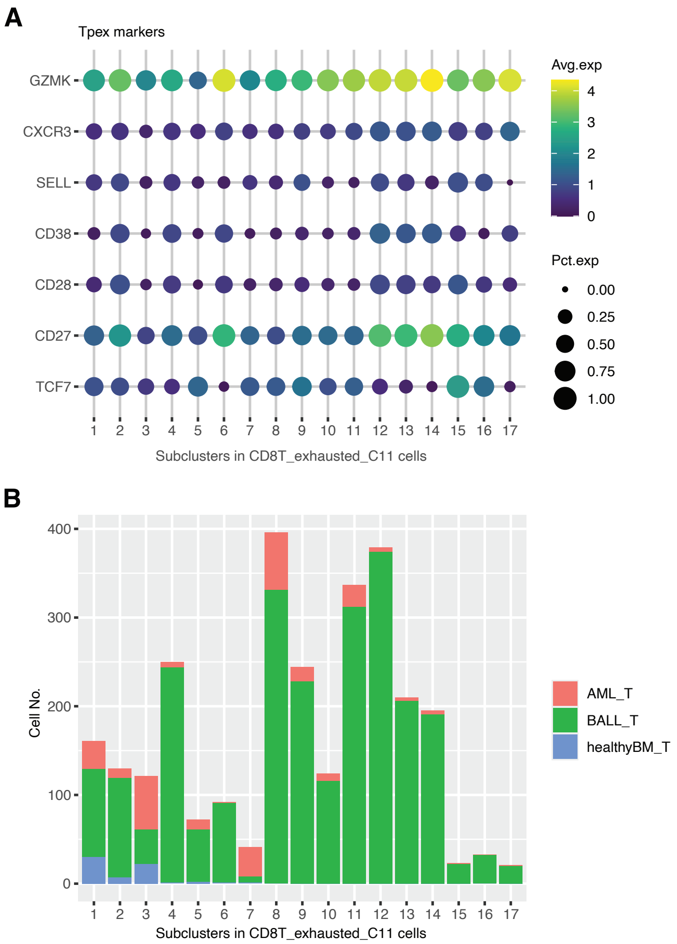


**Supplemental Methods Figure 1: Classification of Tpex and Tex-term cells in CD8T_exhausted_C11.** (**A**) Expression of Tpex marker genes across subclusters. Bubble size indicates the proportion of cells expressing each gene, and color represents the average expression level. (**B**) Number of cells per subcluster in B-ALL, AML, and healthy bone marrow samples.
